# Isolation of folate-producing probiotics and its regulatory effects on homocysteine metabolism and gut microbiota composition

**DOI:** 10.64898/2026.04.29.721560

**Authors:** Mingfang Pan, Changming Ye, Yun Song, Minqing Tian, Runming Wang, Ping Chen

**Affiliations:** Institute of Biopharmaceutical and Health Engineering, Shenzhen International Graduate School, Tsinghua University, Shenzhen, China; Shenzhen Evergreen Medical Institute, Shenzhen, China; Inspection and Testing Center, Key Laboratory of Cancer FSMP for State Market Regulation, Shenzhen 518057, PR China; AUSA Research Institute, Shenzhen AUSA Pharmed Co Ltd, Shenzhen, China; Precision Nutrition Innovation & Transformation Public Service Platform, Shenzhen 518057, PR China

**Keywords:** folate, homocysteine, probiotic, gut microbiota, vitamin B, micronutrient, microbial cross-feeding

## Abstract

**Background:** Folate deficiency is a global nutritional problem associated with multiple adverse health outcomes, including impaired one-carbon metabolism and elevated homocysteine levels (hyperhomocysteinemia). Gut microbiota-mediated folate biosynthesis has emerged as a promising strategy for improving host folate status. This study aimed to isolate folate-producing probiotic strains, clarify their folate synthesis mechanisms, and evaluate their regulatory effects on folate metabolism and gut microbiota in folate-deficient mice.

**Methods:** High-throughput cultivation and screening were performed to isolate folate-producing probiotics. Whole-genome sequencing, pathway reconstruction, and metabolite profiling in fermented milk were used to explore folate biosynthesis pathways and potential microbial cross-feeding interactions. A folate-deficient mouse model was established to evaluate the effects of a probiotic cocktail on serum folate, homocysteine (Hcy) levels, and gut microbiota composition using microbiological assays, biochemical analyses, qPCR, 16S rRNA gene sequencing, alpha diversity analysis, principal coordinates analysis (PCoA), and Linear discriminant analysis Effect Size (LEfSe) analysis.

**Results:** Over 1,000 bacterial isolates were obtained, and over 10 strains, mainly belonging to *Lactobacillus*, *Bifidobacterium*, and *Bacillus*, showed folate production levels above 100 ng/mL. Genomic analysis revealed that most selected probiotic strains lacked genes involved in para-aminobenzoic acid (pABA) biosynthesis but retained downstream folate synthesis modules, suggesting a potential dependence on pABA-producing gut commensals for precursor supply through microbial cross-feeding. In fermented milk, probiotic strains mainly produced bioactive folates (5-MeTHF and THF), with strain-specific production capacities; *L. plantarum*, *W. coagulans*, and *B. animalis* subsp. *lactis* significantly increased 5-MeTHF levels in fermented milk. *In vivo*, high-dose probiotic intervention significantly elevated serum folate (p<0.01) and reduced Hcy (p<0.05) in folate-deficient mice, while medium-dose intervention showed no significant effects. The probiotic strains colonized the mouse gut in a dose-dependent manner: high-dose group exhibited >4,000-fold increase in relative abundance (*Bifidobacteriaceae* and *Bacillaceae* enriched), medium-dose group only enriched *Bacillaceae*, and low-dose group showed no effective colonization. High dose probiotic treatment enhanced gut microbial species diversity (increased Shannon index) and restored folate deficiency-induced gut microbiota dysbiosis (PCoA clustering closer to normal group).

**Conclusion:** This study screened high folate-producing probiotic strains and demonstrated their ability to synthesize active 5-MeTHF, which may rely on microbial cross-feeding in gut microbiota. Furthermore, we demonstrated that folate-producing probiotic intervention significantly improves folate status and Hcy metabolism and restores gut microbiota homeostasis in folate-deficient mice. These findings suggensted that such probiotics could serve as a safer, more physiological intervention for folate deficiency and hyperhomocysteinemia, especially in populations with MTHFR polymorphisms.

## 1. Introduction

Folate is an essential water-soluble B vitamin that plays a pivotal role in one-carbon metabolism, including DNA synthesis, methylation reactions, and amino acid metabolism [1]. Inadequate folate availability is closely associated with elevated plasma homocysteine levels (hyperhomocysteinemia), a recognized risk factor for cardiovascular diseases, neurodegenerative disorders, and metabolic syndrome[2]. Although folic acid supplementation has been widely adopted to prevent folate deficiency, increasing evidence suggests that excessive intake of synthetic folic acid possibly lead to unmetabolized folic acid accumulation and potential adverse health effects, highlighting the need for safer and more physiologically relevant alternatives [3].

Although dietary intake remains the primary determinant of folate and vitamin B12 status, emerging evidence suggests that the gut microbiota may play a contributory role in host folate status[4]. Specific commensal bacteria, particularly lactic acid bacteria and bifidobacteria, intrinsically synthesize biologically active folate derivatives, including 5-methyltetrahydrofolate (5-MeTHF), the predominant circulating form of folate in humans [5, 6]. Unlike synthetic folic acid, which requires enzymatic reduction to become metabolically active 5-MeTHF, bacteria-derived folate is inherently bioavailable and can directly integrate into host metabolic pathways[5, 7]. Thus, folate-producing probiotics represent a promising strategy for improving folate nutrition and regulating homocysteine metabolism through microbiota-mediated mechanisms[8, 9].

Previous studies have reported the isolation of folate-producing probiotics from both traditional fermented foods [10–13] and the human gut [9, 14, 15], yet most of the studies quantify total folate production using microbiological assays or liquid chromatography, lacking subsequent mass spectrometry validation of bioactive folate profiles. This methodological limitation likely reflects the inherent instability of 5-methyltetrahydrofolate (5-MeTHF), the primary bioactive folate,which is readily oxidized into derivative compounds [16]. Furthermore, the therapeutic efficacy of these probiotics (as single strains or consortia) in enhancing host folate bioavailability, modulating homocysteine metabolism, and remodeling gut microbial ecology has not been systematically investigated in animal models or human trials.

In the present study, we isolated and identified folate-producing probiotic strains from traditional fermented foods and human gut microbiota. Their folate-producing capacities were evaluated by quantifying total folate and 5-MeTHF levels using microbiological assays and LC-MS/MS analysis, respectively. Furthermore, probiotic cocktails composed of selected high-producing strains were administered to mice and human subjects to investigate their effects on homocysteine metabolism and gut microbiota composition. This work provides new insights into the functional role of folate-producing probiotics as a natural and microbiota-targeted approach for modulating host homocysteine metabolism and improving metabolic health.

## 2. Materials and Methods

### 2.1 Isolation and Identification of Probiotic Strains

Potential folate-producing were isolated from Tibet kefir and human stool by an established high-throughput workflow adapted from previously reported works [11, 17]. Briefly, ∼1 g sample was suspended in 10 ml 1×PBS and a series of dilutions were prepared and spread culture on the MRS agar plates, mixed with 2% CaCO_3_, anaerobically incubating for 48 h at 37 ◦C. The bacterial colonies which formed clear zones around it by dissolving CaCO_3_ were isolated at random. Then each isolated strain was anaerobically incubated MRS broth at 37 ◦C for 24 h for without shaking. Different culture cells were collected, washed, and resuspended with sterilized1×PBS. Subsequently, the obtained suspensions were inoculated into Folic Acid Assay Broth (BioTAI, Beijing, China) which free of folic acid and anaerobically incubated at 37 ◦C for 48 h. The growth was determined 600 nm using a Spectra microplatereader (Multiskan FC, thermo scientific). Strains with OD_600_ above 0.2 were deemed to have successful bacterial growth and were identified as potential folate producers.

### 2.2 Evaluation of Folate-Producing Ability

Potential folate producers were further incubated in MRS broth anaerobically for 24 h and the bacterial cultures were collected and centrifuged at 5,000 × g for 15 min at 4◦C. Bacterial culture supernatants were reserved for folate quantification by microbiological methods as previously reported [16]. Total folate was quantified by microbiological methods as previously described. Briefly, 50 uL of bacterial culture was filtered by 0.2 uM filter and then added to 96-well plate ( BioTAI, Beijing, China) loaded with Folic Acid Assay Broth. The plate was internally coated coated with freeze-dry *Lactobacillus rhamnosus* ATCC 12126, an auxotrophic strain for folate. Various concentrations of folate standard ranging from 16 ng/mL to 128 ng/mL were prepared for standardization. After cultured in the dark for 48h,the growth was determined by measured the 600 nm. The concentration of the bacterial culture was calculated against standards concentration. The bacterial cultures that exceed the standard range were further diluted accordingly and measured again.

The isolates with efficient folate-producing ability was further identified by 16S ribosomal DNA (rDNA) gene sequencing, and constructing the phylogenetic tree using the MEGA 5.1 package [18].

### 2.3 Determination of Folate and folate-derived metabolits in fermented milk by LC-MS/MS

LC-MS/MS was employed to determine folate and its derived metabolites in fermented milk as previously reported with modification [12, 16, 19]. Briefly, the samples were initially spiked with three mixed internal standard solution([13C5]-Folic Acid (SHANGHAI ZZBIO CO., LTD), deuterated labeled Folinic acid(Toronto Research Chemicals), calcium d,l-5-methyltetrahydrofolate(SHANGHAI ZZBIO CO., LTD) and treated with enzymes including papain, α-amylase and rat serum. After enzymatic hydrolysis, the mixtures were centrifugated at 10,000 r/min and 4 °C for 5 min. The supernatant was methanol precipitated, filtered and dry via nitrogen-blow and reconstituted for analysis by LC-MS/MS.

The analysis was conducted using an LC-MS/MS system with a KineteX C18 column (2.6 μm, 2.6 × 3 mm, MS-062) and a 4500 triple quadrupole mass spectrometer (AB SCIEX) equipped with an ESI source. The chromatographic conditions were as follows: column temperature 40 °C, auto-sampler temperature 4 °C, flow rate 0.4 mL/min, injection volume 10 μL, and mobile phase consisting of 0.1% formic acid in water (A) and 0.1% formic acid in methanol (B) with a gradient elution program. The mass spectrometer was operated in positive ion mode with MRM scanning, and the key parameters included collision gas (CAD) 9, curtain gas (CUR) 25, nebulizer gas (Gas1) 60.0, auxiliary gas (Gas2) 60.0, spray current (IS) 5500 V, and atomization temperature (TEM) 550 °C.

Standard curves were prepared for quantification: mixed folate standard solutions (folate, 6S-5-methyltetrahydrofolate, 5-formyltetrahydrofolate)were serially diluted to prepare standards. Additionally, standard curves for N-(4-aminobenzoyl)-L-glutamic acid, Mefox, and tetrahydrofolate were prepared through stepwise dilution. All the standards were from Sigma-Aldrich. Quality control (QC) solutions were prepared using QC reserve solutions to ensure analytical reliability.

### 2.4 Animal grouping and treatment

All animal procedures were performed in accordance with the Guidelines for Care and Use of Kezhuo Medical Laboratory Testing Co., Ltd (Shenzhen, China) and approved by the Animal Ethics Committee of Kezhuo Medical Laboratory Testing Co., Ltd (Shenzhen, China). Male C57BL/6 mice (5–6 weeks old, ∼18 g; n = 36) were used in this study. After one week of acclimatization, the mice were randomly assigned to a normal diet group (Nd, n = 6) or a folate-deficient diet group (n = 30). Mice in the Nd group were fed a standard chow diet, whereas mice in the folate-deficient group were fed a folate-deficient diet (AIN-93; Ready Dietech). After one month of dietary intervention, 10 μL of blood was collected from the tail vein, and serum folate concentrations were measured using a microbiological assay to confirm successful establishment of the folate-deficient mouse model. Subsequently, mice were subjected to intervention with a probiotic cocktail consisting of seven isolated probiotic strains: *Lactiplantibacillus plantarum* FL001, *Lactiplantibacillus plantarum* Ausa002, *Lactiplantibacillus plantarum* Ausa004, *Bifidobacterium longum* subsp. longum FL006, *Bifidobacterium breve* FL008, *Weizmannia coagulans* AUSA001, and *Bifidobacterium animalis* subsp. lactis BB100.

Mice fed the folate-deficient diet were further subdivided into the following groups: folate-deficient diet control (FA_d), high-dose probiotic cocktail (Pro_H), medium-dose probiotic cocktail (Pro_M), low-dose probiotic cocktail (Pro_L), and synthetic folic acid positive control (FA). Mice in the Nd and FA_d groups were orally gavaged with 200 μL/day of sterile 1× PBS. Mice in the Pro_H group received 200 μL/day of probiotic cocktail suspension at a concentration of 1 × 10¹⁰ CFU/mL, whereas mice in the Pro_M and Pro_L groups received probiotic suspensions at concentrations of 1 × 10⁹ CFU/mL and 1 × 10⁸ CFU/mL, respectively. Mice in the FA group were orally gavaged with 200 μL/day of folic acid solution (9.125 μg/mL), corresponding to a dose of 73 μg/kg body weight.

After 6 weeks of probiotic intervention, fecal and blood samples were collected. Quantitative PCR (qPCR) using primers specific to the administered probiotic species was performed to evaluate probiotic strain colonization. Serum folate and homocysteine (Hcy) levels were determined using chemiluminescent immunoassays. The experimental design is illustrated in Figure 3A.

#### Quantitative real-time PCR (qPCR) detection of probiotic species

Quantitative real-time PCR (qPCR) was performed to monitor the changes in the corresponding probiotic species in the mice feces following oral gavage. Species-specific primers targeted the bacterial strains administered were adapted from previously publications (Supplementary Table 1). The relative changes in bacterial abundance were calculated using the 2^^ΔΔCt^ method.

#### Gut microbiota sequencing and bioinformatics

Extraction of fecal DNA and sequencing were performed using previously described methods [17]. Briefly, bacterial genomic DNA was extracted from fecal samples using MagPure Stool DNA KF Kit B (MAGEN, GuangZhou, China) according to manufacturers’ protocols. The extracted DNA was used as the template to amplify V3-V4 variable region of 16S rDNA of bacteria by forward and reverse PCR degenerate primers F and R (338F: ACTCCTACGGGAGGCAGCAG, 806R: GGACTACHVGGGTWTCTAAT). Sequencing reads of PE300 bases length are generated with DNBSEQ-G400 platform (BGI-Shenzhen, China). Amplicon Sequence Variants (ASVs) sequences are generated by clustering denoised sequences with 100% similarity. Operational taxonomic units (OTUs) with higher than 97% similarity were delineated using the USEARCH (v7.0.1090)[20]. OTU representative sequences are aligned against the database for taxonomic annotation by RDP classifer (v2.2) software[21]. Alpha diversity is mainly measured by shannon index and chao index using mothur(v.1.31.2) [22]. Community dissimilarity was analyzed by Principal component analysis (PCA). Weighted Unifrac distance between each group was measured, and pairwise Permutational Multivariate Analysis of Variance (PERMANOVA) test was used to test the significant difference between each group. The software used in this step was the ‘ade4’ package in R software (v3.1.1) [23].

### 2.6 Statistical Analysis

All statistical analyses were performed using GraphPad Prism 9 (GraphPad Software, Inc.). Statistical differences between experimental groups were evaluated by Student’s t tests and one-way analysis of variance (ANOVA) with a Dunnett’s test for multiple comparisons. All data were expressed as means ± standard deviations (SD). P values of <0.05 were considered statistically significant.

## 3. Results

### 3.1 Isolation of folate-producing probiotic strains

A total of more than 1,000 bacterial isolates were obtained through high-throughput cultivation and screening. Quantitative analysis by microbiological method identified over 10 strains capable of producing at levels exceeding 100 ng/mL folate, primarily belonging to the genera *Lactobacillus*, *Bifidobacterium*, and *Bacillus* (Table 1, Supplementary Figure 1). These strains were selected for further genomic and functional analysis.

**Table 1.** Total folate production of the isolated probiotic strains quantified by microbiological assay.

| No. | Isolated probiotic strains | Sources | Supernatant folate (ng/ml) |
| --- | --- | --- | --- |
| 1 | *Bacillus cereus* AF003 | Fermented food | 98.7 ± 7.8 |
| 2 | *Bacillus cereus* AUSA008 | Human stool | 129.5 ± 11.5 |
| 3 | *Streptococcus thermophilus* AF006 | Fermented food | 127.0 ± 20.8 |
| 4 | *Streptococcus thermophilus* AF007 | Fermented food | 84.3 ± 13.6 |
| 5 | *Lactocaseibacillus paracasei* AF008 | Fermented food | 0.6 ± 0.2 |
| 6 | *Bifidobacterium breve* FL008 | Human stool | 62.7 ± 19.9 |
| 7 | *Bifidobacterium breve* AUSA012 | Human stool | 51.2 ± 24.0 |
| 8 | *Bifidobacterium adolescentis* AUSA013 | Human stool | 62.7 ± 4.4 |
| 9 | *Bifidobacterium longum* subsp. longum AUSA015 | Human stool | 136.7 ± 15.1 |
| 10 | *Bifidobacterium longum* subsp. longum AUSA016 | Human stool | 72.1 ± 15.0 |
| 11 | *Bifidobacterium longum* subsp. longum FL006 | Human stool | 114.0 ± 24.0 |
| 12 | *Lactiplantibacillus plantarum* AUSA002 | Fermented food | 113.4 ± 13.2 |
| 13 | *Lactiplantibacillus plantarum* AUSA004 | Human stool | 113.0 ± 12.8 |
| 14 | *Lactiplantibacillus plantarum* FL001 | Human stool | 131.0 ± 19.7 |
| 15 | *Lactiplantibacillus plantarum* AUSA019 | Human stool | 89.6 ± 24.7 |
| 16 | *Lactocaseibacillus rhamnosus* AF009 | Human stool | 1.4 ± 0.6 |
| 17 | *Lactocaseibacillus rhamnosus* AUSA011 | Human stool | 0.8 ± 0.2 |
| 18 | *Lactocaseibacillus rhamnosus* AUSA020 | Fermented food | 0.7 ± 0.5 |
| 19 | *Limosilactobacillus fermentum* AOSA CECT100 | Fermented food | 2.5 ± 1.3 |
| 20 | *Bifidobacterium animalis* subsp. lactis AOSA BB100 | Fermented food | 3.6 ± 1.7 |
| 21 | *Bifidobacterium animalis* subsp. lactis AOSA BB100A | Fermented food | 3.6 ± 1.1 |
| 22 | *Bifidobacterium animalis* subsp. lactis AOSA BB100B | Fermented food | $2.5 \pm 1.0$ |
| 23 | *Bifidobacterium animalis* subsp. lactis AOSA BB100C | Fermented food | $5.8 \pm 1.8$ |
| 24 | *Bifidobacterium bifidum* AOSA R100 | Human stool | $68.6 \pm 16.9$ |
| 25 | *Lactobacillus acidophilus* AOSA NCFM100 | Fermented food | $1.1 \pm 0.4$ |
| 26 | *Lactobacillus acidophilus* AOSA NCFM100A | Fermented food | $1.1 \pm 0.3$ |
| 27 | *Weizmannia coagulans* FL009 | Fermented food | $79.6 \pm 12.2$ |
| 28 | *Weizmannia coagulans* FL010 | Fermented food | $53.9 \pm 9.2$ |
| 29 | *Weizmannia coagulans* FL011 | Fermented food | $43.4 \pm 20.5$ |
| 30 | *Weizmannia coagulans* AUSA001 | Human stool | $114.4 \pm 26.8$ |
| 31 | *Lactobacillus delbrueckii* subsp. bulgaricus FL012 | Fermented food | $1.9 \pm 0.5$ |

### 3.2 Strain-specific folate biosynthesis pathways and evidence of microbial cross-feeding

Whole-genome sequencing and pathway reconstruction of 34 commonly used probiotic strains revealed substantial heterogeneity in folate biosynthesis gene content among these strains (Figure 1). Most strains lacked genes involved in para-aminobenzoic acid (pABA) biosynthesis, the key folate precursors, while retaining partial downstream folate biosynthesis modules (Figure 1A). These results suggest that folate production by these probiotics is genetically incomplete and potentially dependent on external precursor supply.

**Figure 1.**
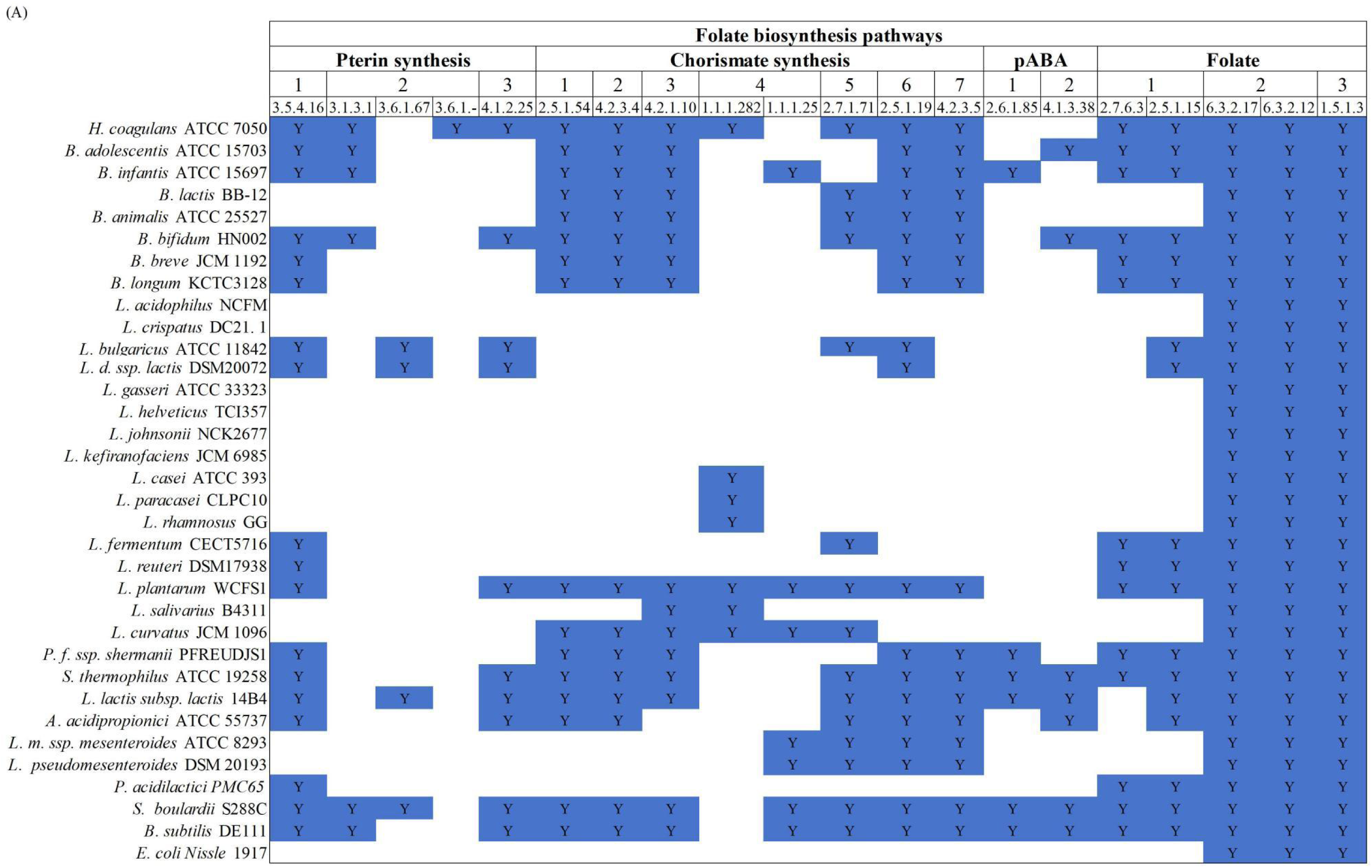
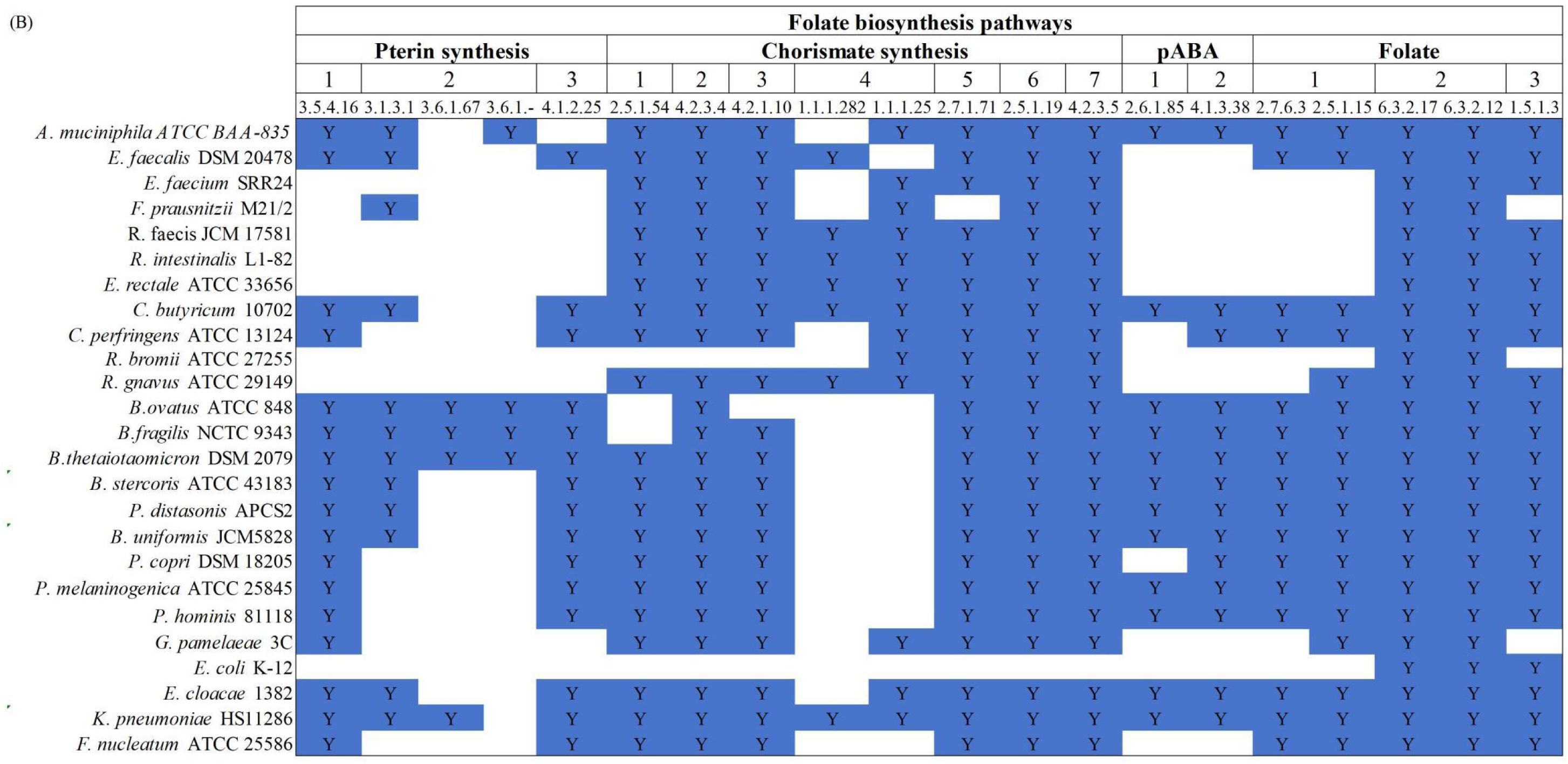
Folate denovo synthesis pathways analysis in (A) probioitc strains and (B) common gut commensals. Y: presence of the corresponding gene. The protein sequences the probiotic strains and gut commensals were downloaded from NCBI database (https://www.ncbi.nlm.nih.gov/). KofamKOALA (https://www.genome.jp/tools/kofamkoala/) was used to assign K numbers to the sequences by HMMER/HMMSEARCH against KOfam (a customized HMM database of KEGG Orthologs (KOs)). The predicted KOs ssociated with chorismate synthesis, pABA synthesis, pterin synthesis, and folate synthesis were mapped to folate biosynthesis pathways by KEGG maper (https://www.kegg.jp/kegg/mapper)

In contrast, genomic analysis of 25 representative gut commensals from six phyla (Firmicutes, Bacteroidetes, Actinobacteria, Proteobacteria, Verrucomicrobia, Fusobacteria) showed that several commensals, particularly members of *Bacteroides* and *Prevotella*, harbored pABA biosynthesis-related genes, suggesting functional complementarity within the gut ecosystem.

### 3.3 Characterization of folate and folate-derived metabolite production in probiotic-fermented milk

Analysis of folate and folate-derived metabolites in probiotic-fermented milk showed that the main folate forms produced by the probiotic strains were the biologically active vitamers 5-methyltetrahydrofolate (5-MeTHF) and tetrahydrofolate (THF), while folate-producing capacity varied considerably among strains (Figure 2). The *L. plantarum* species, *W. coagulans* species, *B. animalis* subsp. *lactis*, and the fermentation starter (*S. thermophilus* and *L. delbrueckii* subsp. *bulgaricus*) significantly increase the 5-MeTHF amount in fermented milk. Only *L. plantarum* species and *W. coagulans* species significantly increase the THFA concentration. In contrast, it showed the based amount of 5-MeTHF in the fermented milk (with starter) decreased after the addition with *L. rhamnosus*. This consistent with the folate synthesis pathway analysis which showed *L. rhamnosus* lack the denovo folate synthesis pathway and has to scavage folate from external environment. Interestingly, *B. animalis* subsp. *lactis* genome analysis showed lack of denovo folate systhesis pathway but showed relatively high ability to produce 5-MeTHF, indicated that the milk fermented with starter contain the percursor of folate denovo synthesis.

**Figure 2.**
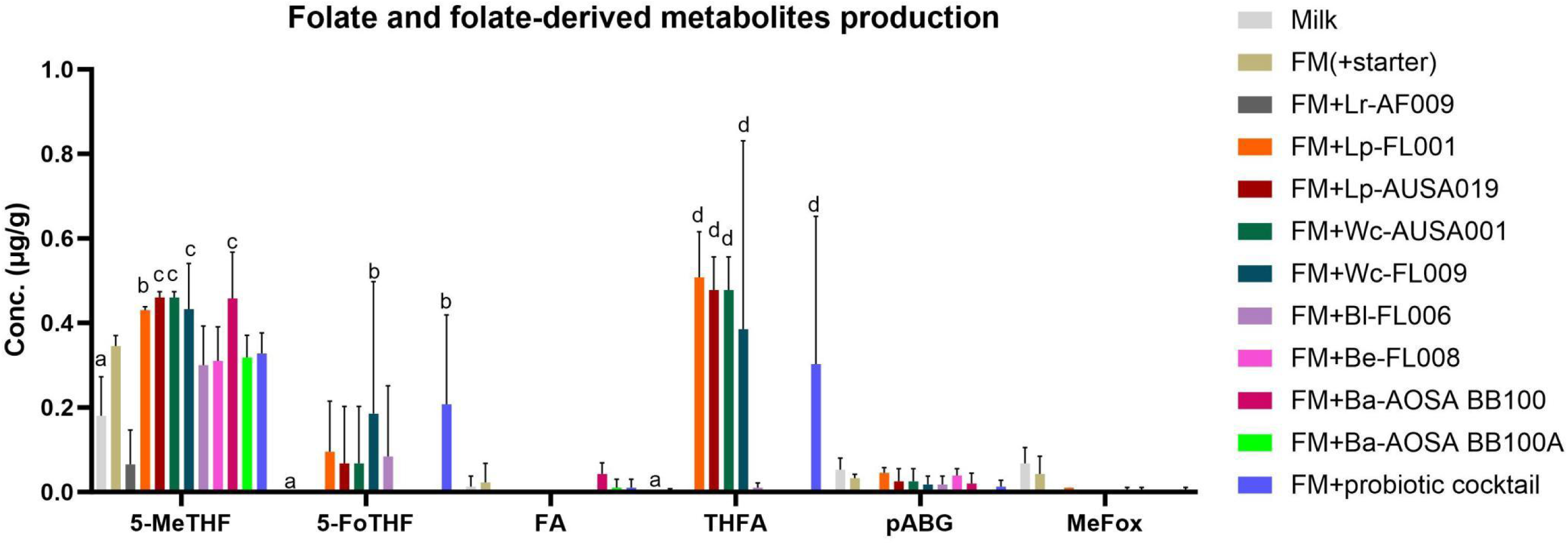
Folate and folate-derived metabolites production in milk fermented by probiotic strains. Milk was fermented with starter (*Streptococcus thermophilus* AF006 and *Lactobacillus delbrueckii subsp. bulgaricus* FL006) and probiotic strains for 18 h, folate and folate-derived metabolites produced were determined by LC-MS/MS. 5-MeTHF:5-methyltetrahydrofolate, 5-FoTHF:5-formyltetrahydrofolate, FA:folic acid, THF:tetrahydrofolate, pABG:p-aminobenzoylglutamate, MeFox: (6RS)-MeFox.

### 3.4 Effects of probiotic cocktail on mice folate and homocysteine metabolism

Folate-deficient diet was used to establish a folate-deficient mice model (Figure 3A). Mice body weight was record during the intervention. Neither folate-deficient diet nor probiotic intervention significantly affect the body weight (Supplementary Figure 2).

**Figure 3.**
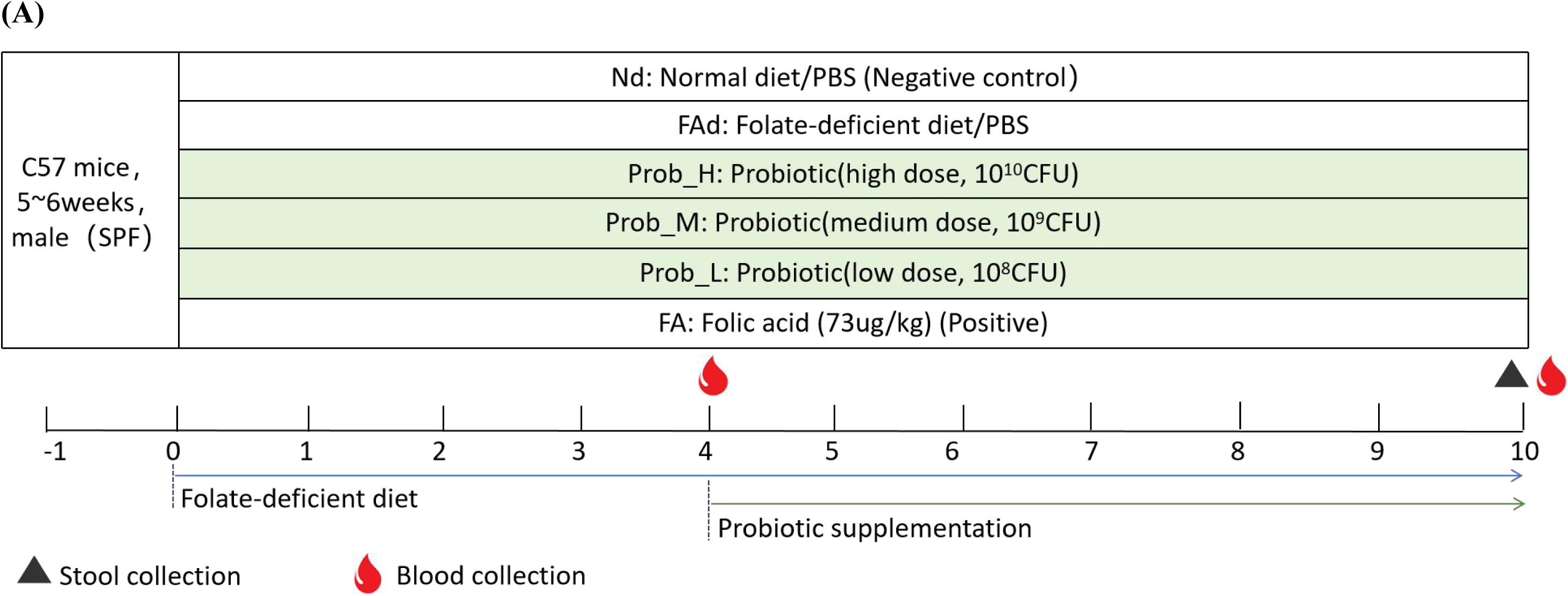
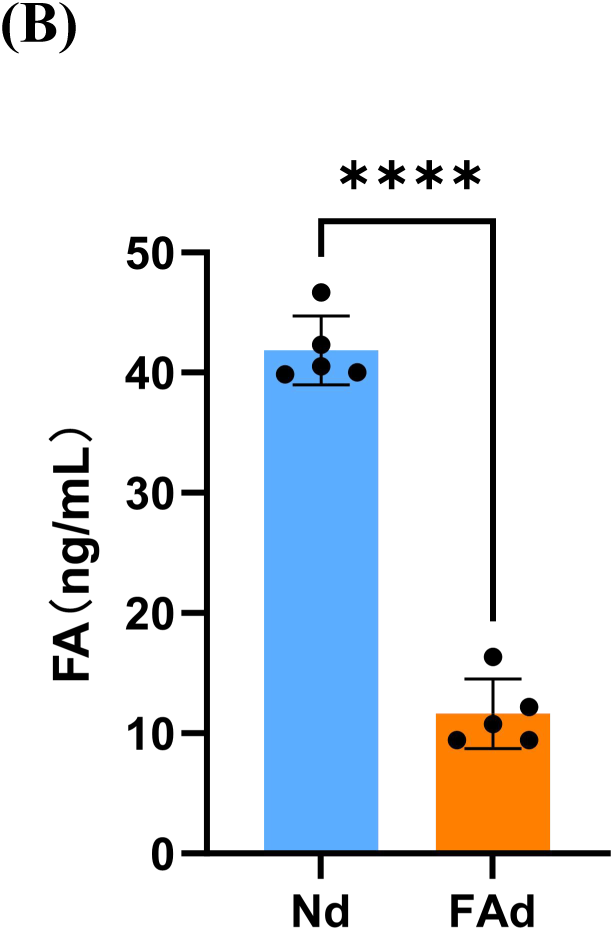
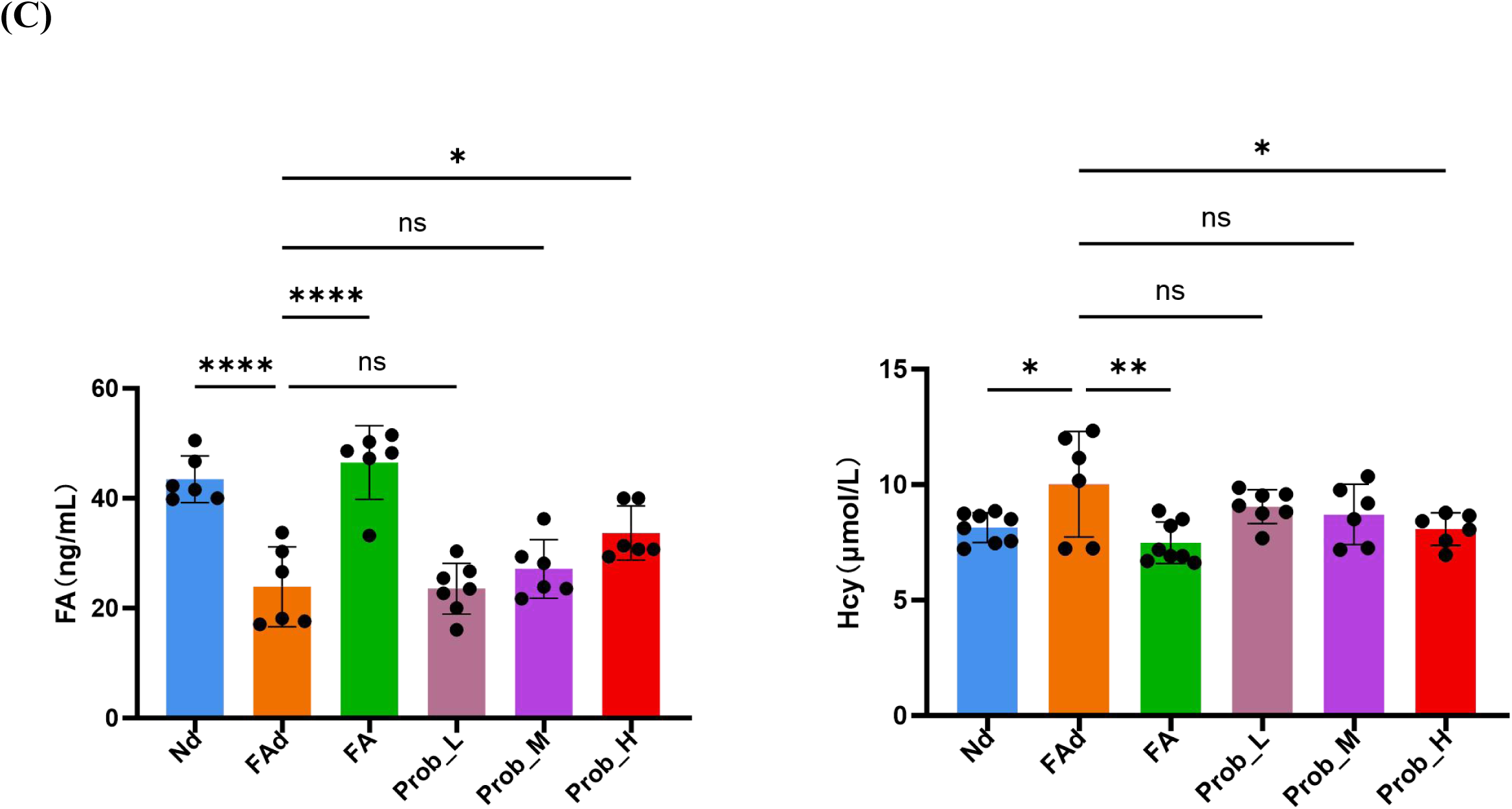
Probiotic cocktail administration improves folate status and reduces homocysteine levels in folate-deficient mice. (A) mice group and intervention design. Nd: Normal diet/PBS (Negative control); FAd: Folate-deficient diet/PBS; Prob_H: Probiotic (high dose, 10^10^CFU); Prob_M: Probiotic(medium dose, 10^9^CFU); Prob_L: Probiotic (low dose, 10^8^CFU); FA: Folic acid (73ug/kg) (Positive). (B) Mice folate level determined by microbiological assays after 1 month folate-decifient diet. (C) mice folate and homocysteine level determined by chemiluminescence assays after 6 weeks probiotic cocktail administration. Data are presented as mean ± SD. Statistical significance was determined by xxx

Following one month feeding with a folate-deficient diet, blood was collected from the tail vein of five mice. Serum folate levels were determined using a microbiological assay. The results demonstrated that serum folate levels were significantly reduced in the folate-deficient group compared to the normal diet group (Figure 3B), indicating the successful establishment of a folate-deficiency mouse model after one month of dietary intervention.

After 6 weeks of probiotic intervention, blood was collected from the orbital sinus. Serum folate and homocysteine (Hcy) levels were quantified using biochemical assays. The results showed that serum folate levels were significantly elevated in the high-dose probiotic group (p<0.01), while a non-significant increasing trend was observed in the medium-dose group (p>0.05) (Figure 3C). Correspondingly, serum Hcy levels were significantly increased in the folate-deficient diet group. Administration of synthetic folate significantly decreased serum Hcy levels (p<0.01). Similarly, a significant reduction in serum Hcy was observed in the high-dose probiotic group (p<0.05), whereas no significant changes were detected in the medium- and low-dose groups.

### 3.5 The colonization and survival ability of probiotics in mice

The qPCR results demonstrated that the probiotic strains increased in a dose-dependent manner in the mice gut after administration (Figure 4). The most significant increase in relative abundance was observed in the high-dose group, with a maximum increase of more than 4000-fold. In contrast, the increase in the low-dose group was limited to less than 100-fold. These results confirmed the presence of administered probiotic strains in fecal samples following oral gavage of high-dose probiotic cocktail, indicating successful colonization of the administrated probiotic in the mice gut (Figure 4). At the strain level, differences in intestinal persistence among the administered strains were also observed. In the high-dose group, *Bifidobacterium* was detected, while in the medium-dose group, only *B. coagulans* was identified. *L. plantarum* was not detected in any of the dose groups.

**Figure 4.**
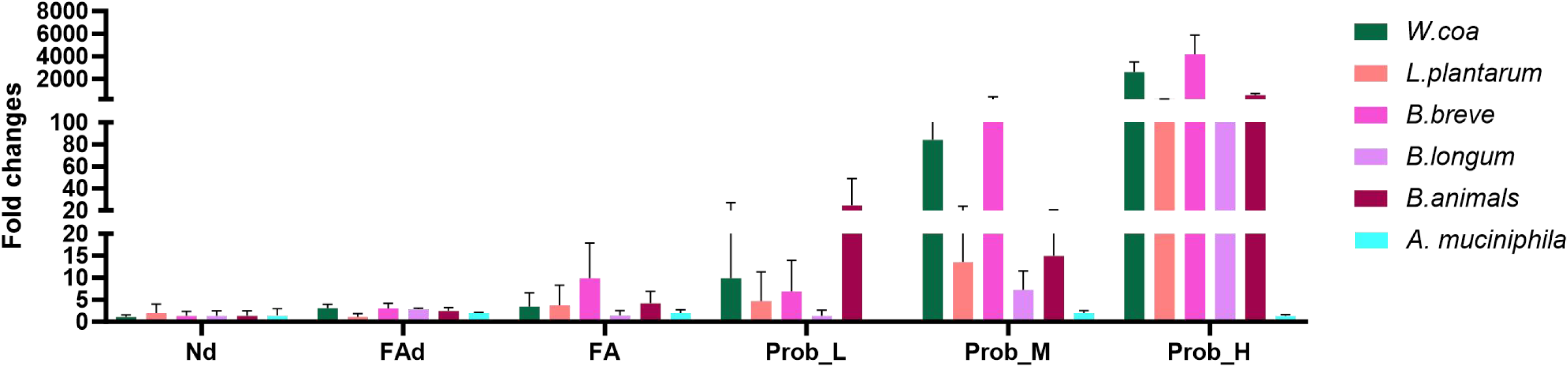
qPCR analysis of the relative abundance of probiotic strains in mouse feces after probiotic cocktail administration. The relative abundance of the administered strains, including *Lactiplantibacillus plantarum*,, *Bifidobacterium longum* subsp. longum FL006, *Bifidobacterium breve* FL008, *Weizmannia coagulans* AUSA001, and *Bifidobacterium animalis* subsp. lactis BB100) in mice feces was quantified by qPCR. . The relative changes in bacterial abundance were calculated using the 2^ΔΔCt method.

### 3.6 Effects on mice gut microbiota composition

Analysis of alpha diversity indices revealed no significant differences in the Shannon and Chao1 indices among the groups following feeding with a folate-deficient diet or probiotic treatment, except for a significant increase in the Shannon index in the high-dose probiotic group compared to the control group, indicating that gavage with a high-dose probiotics was beneficial for increasing gut microbial species diversity (Figure 5A).

**Figure 5.**
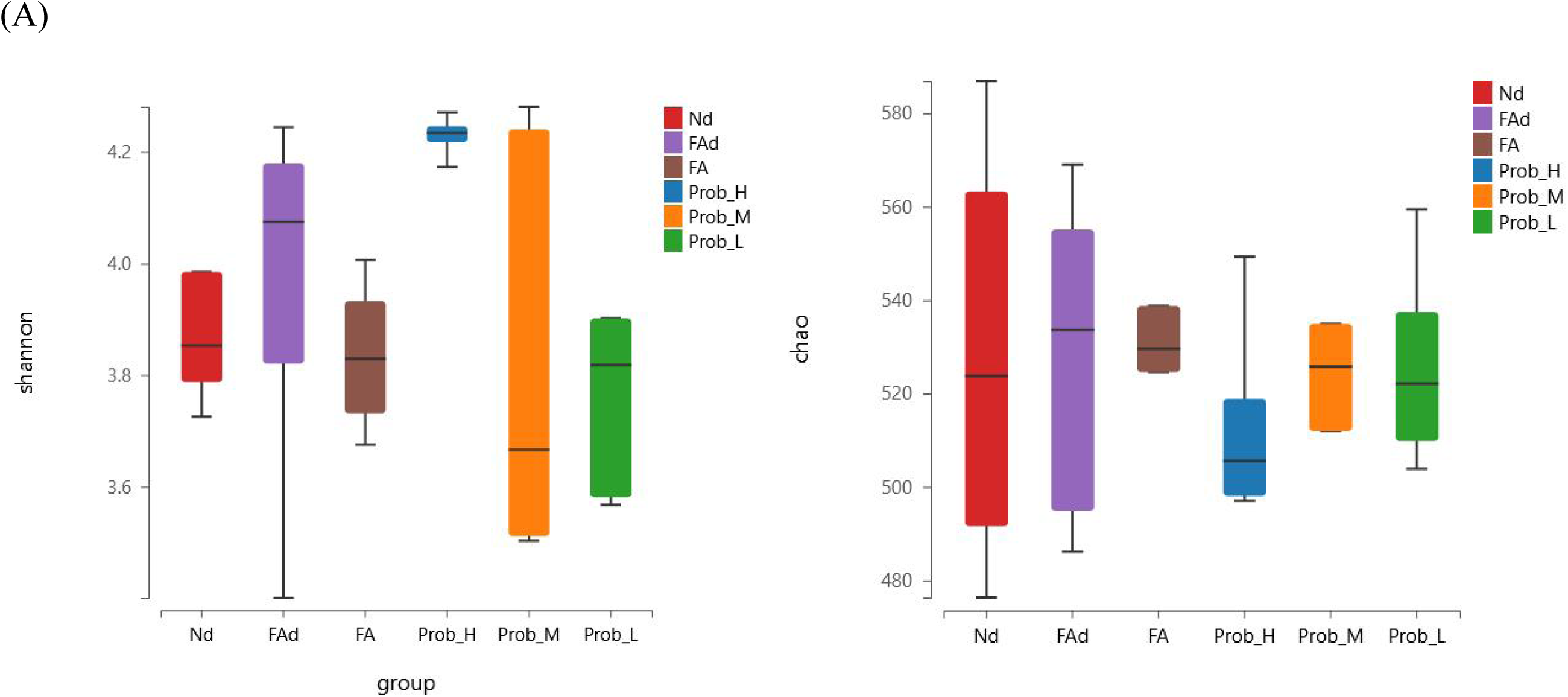
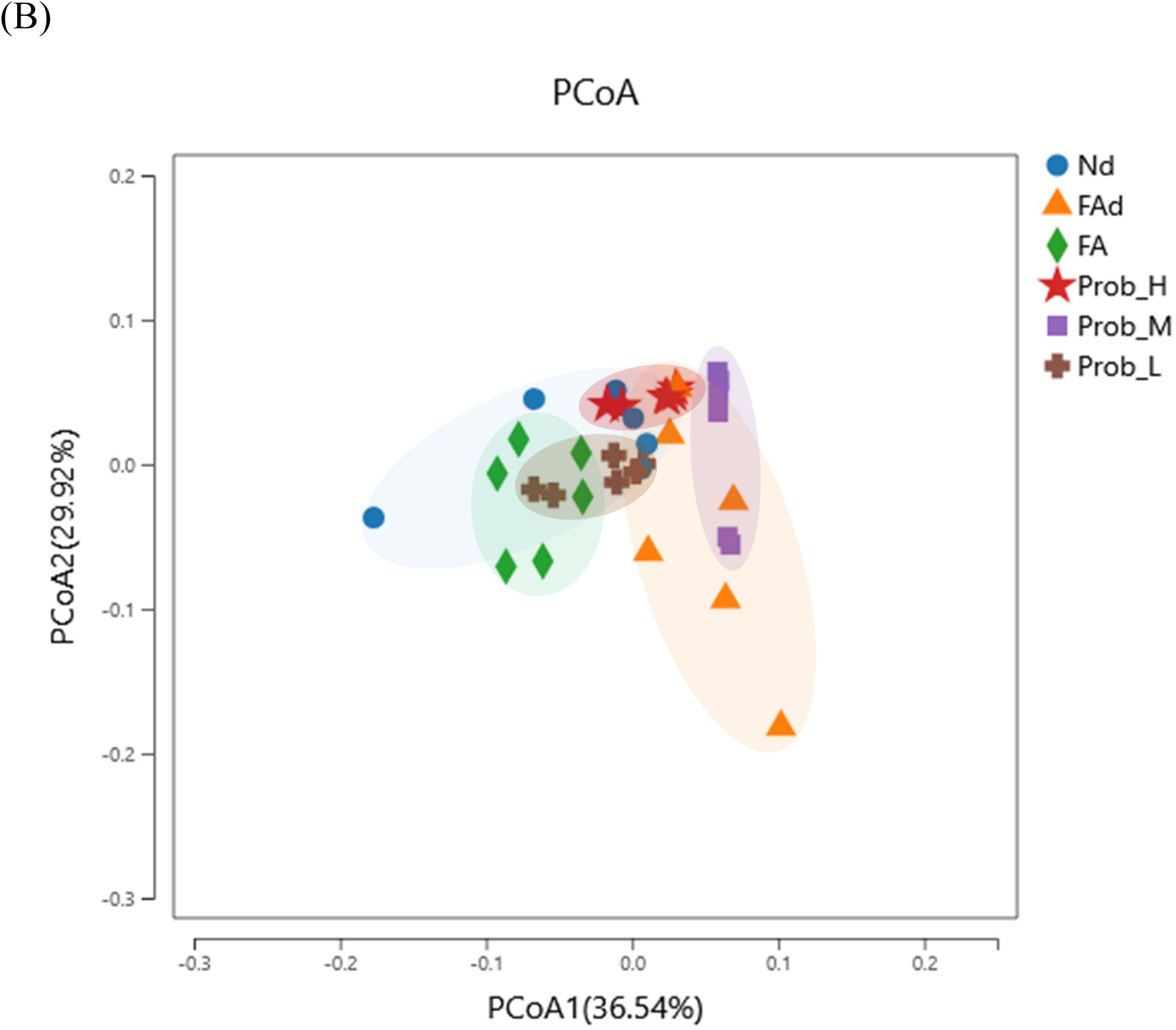
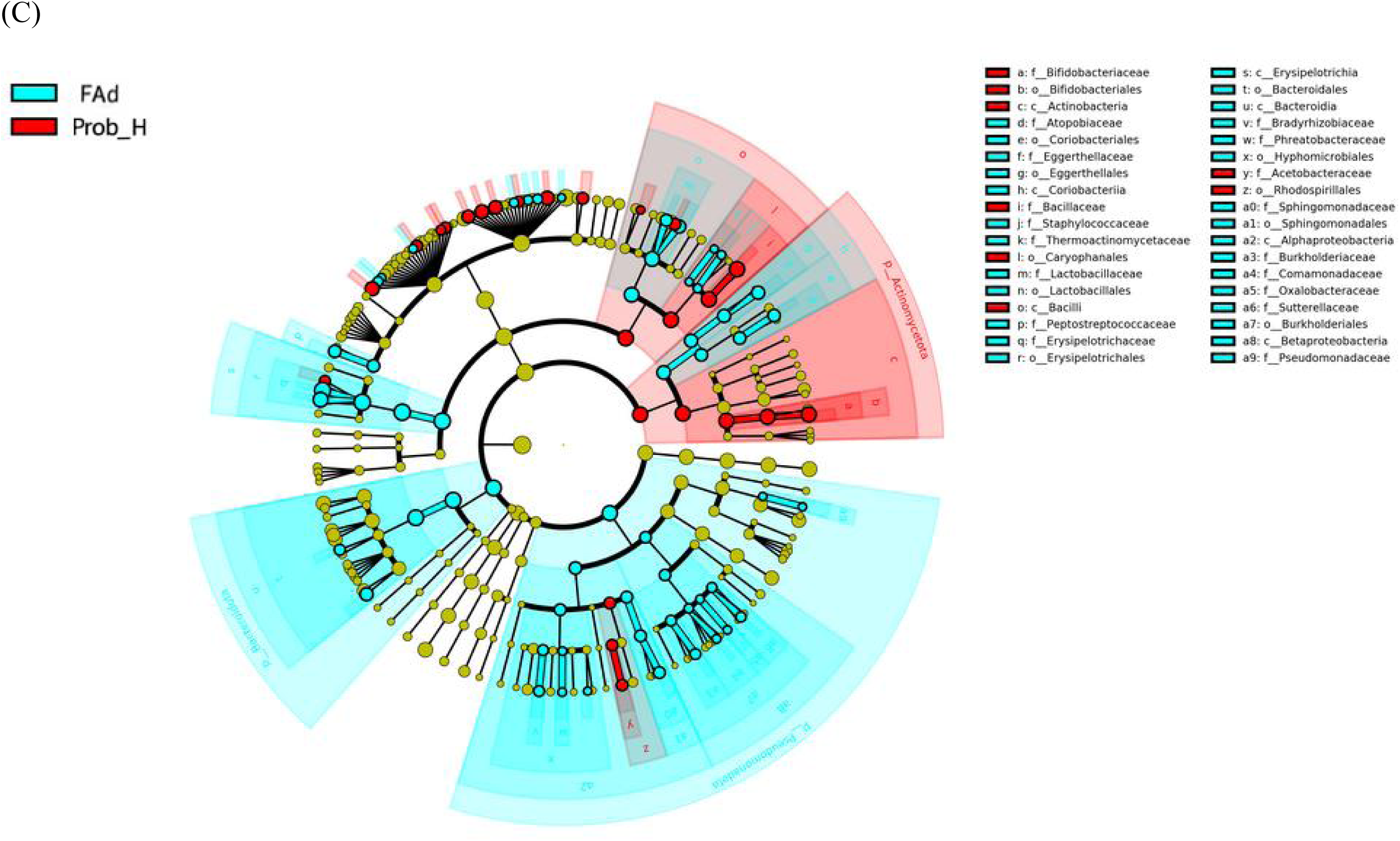
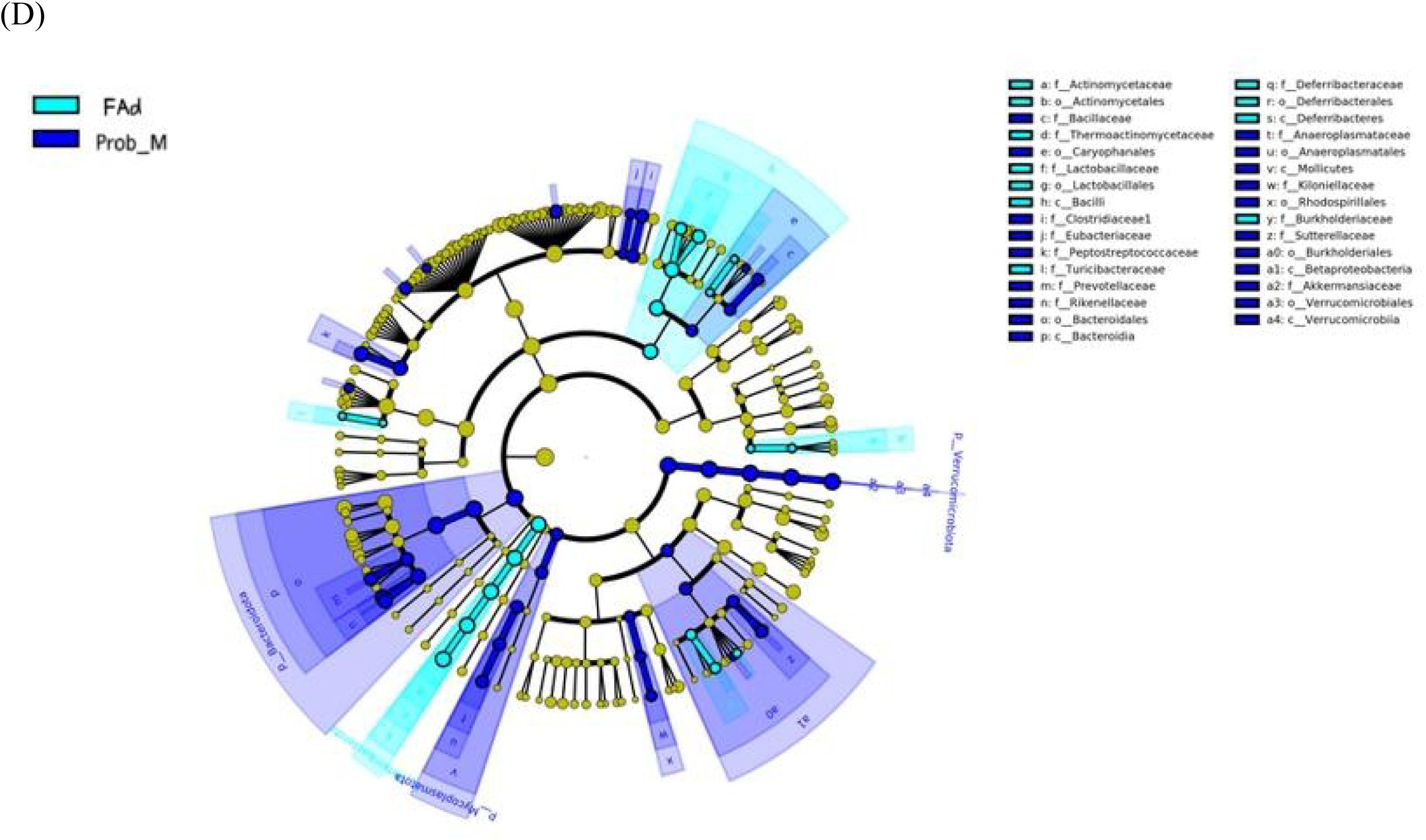
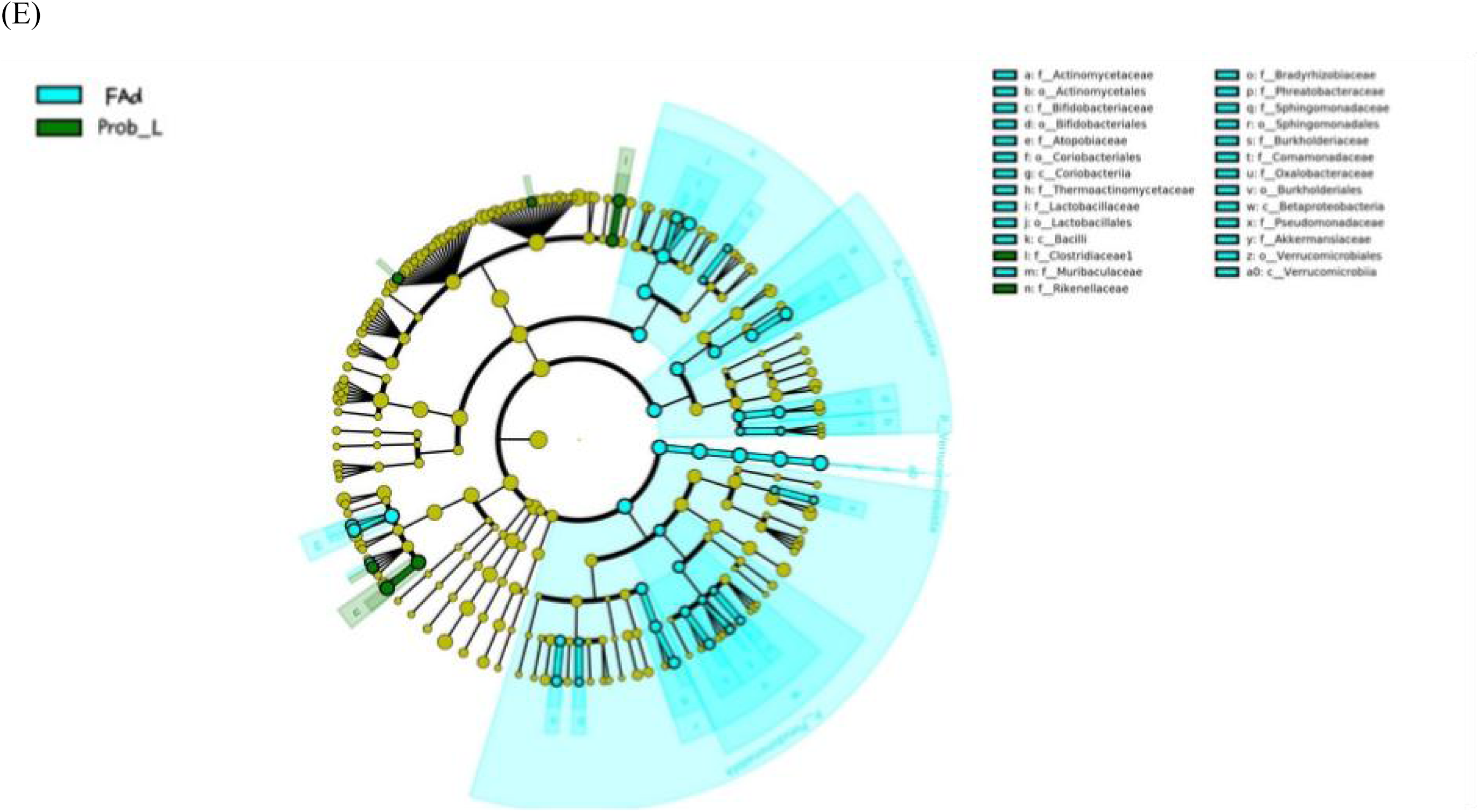
Effects of the probiotic cocktail on gut microbiota composition in mice. (A) Comparison of alpha diversity (Shannon index and Chao index) of the gut microbiota based on groups. (B) Principal coordinates analysis (PCoA) of the gut microbiota based on groups. (C, D, E) Differentially abundant bacterial taxa among groups.

PCoA analysis of the overall microbial composition showed that the gut microbiota of mice fed a folate-deficient diet exhibited a global shift compared to the normal diet group (Figure 5B). However, supplementation with probiotics or synthetic folate shifted the microbial composition closer to that of the normal group, suggesting a tendency of the gut microbiota to return to a normal state following these treatments.

LEfSe analysis was performed to identify differentially abundant bacterial taxa among the groups. As shown in Figure 4C, mice receiving the high-dose probiotic were significantly enriched in Bifidobacteriaceae (order Bifidobacteriales) and Bacillaceae (class Bacilli), indicating successful colonization and proliferation of the administered probiotics in the gut. In the medium-dose group (Figure 4D), enrichment was observed only for Bacillaceae. No enrichment of the administered probiotic strains was detected in the low-dose group (Figure E). These results were consistent with the qPCR data, suggesting that the probiotic strains administered at high and medium doses were able to survive in the mouse gut, whereas those administered at the low dose were not.

## Discussion

This study demonstrates that selectively screened folate-producing probiotic strains can synthesize bioactive folate, particularly 5-methyltetrahydrofolate (5-MeTHF), and supplementation with these strains improves host folate status and homocysteine (Hcy) metabolism in vivo. These findings provide both mechanistic and functional evidence supporting the role of probiotics in micronutrient status and Hcy metabolism.

We characterized the folate-producing capacity of the isolated strains by measuring the total folate by microbiological methods. Consistent with previous studies [9, 11, 13, 15], isolated probiotic strains from representing bacterial species produced variable amount of folate in MRS broth. Genome-based folate pathway reconstruction, however, revealled that *de novo* folate biosynthesis is generally incomplete across probiotic strains. A common limitation was the absence of genes required for para-aminobenzoic acid (pABA) synthesis, a key precursor for folate production. This metabolic deficiency indicated that these probiotic strains rely on exogenous precursors supplied by the MRS broth to support folate production. This observation may further account for the discrepancies in folate levels reported across distinct studies, as the composition of culture media employed may vary between investigations [24].

In contrast, many prevalent gut commensals, e.g. Bacteroides and Prevotella, encoded more complete folate biosynthesis pathways, including pABA-related functions. This complementarity supports a potential cross-feeding model in which probiotics may rely on microbiota-derived precursors (e.g., pABA or related intermediates) to support folate formation in the gut. Such metabolic cooperation is consistent with the ecological organization of the intestinal community, where nutrient exchange can enable functional outputs that are not predictable from the metabolic capacity of individual species alone [25].

Further, we characterized the molecular forms of folate produced by screened probiotic using LC-MS/MS. The results confirmed that the folate produced by isolated probiotic strains in fermented milk are largely the physiologically active folate 5-methyltetrahydrofolate (5-MeTHF). Interestingly, genome analysis revealled *B. animalis* species encoded only a downstream segment of the folate biosynthesis pathway (from dihydropteroate to THF-polyglutamate; including enzymes annotated as EC 6.3.2.17, EC 6.3.2.12, and EC 1.5.1.3). However, isolated *B. animalis* AOSA BB100 generated substantial amounts of 5-MeTHF and THF during milk fermentation. This discrepancy highlights two key points: (i) the presence/absence of canonical genes does not necessarily translate into functional folate production under specific growth conditions, and (ii) standard laboratory media may not reflect the precursor availability, redox state, or metabolic cues present in food matrices or the gut. Thus, functional validation under biologically relevant conditions remains essential alongside in silico prediction.

5-MeTHF is the predominant bioactive form circulating in humans and can be utilized directly in one-carbon metabolism without requiring extensive enzymatic conversion [26]. In contrast, synthetic folic acid must first be reduced and converted into 5-MeTHF, a process that depends in part on methylenetetrahydrofolate reductase (MTHFR) activity. Reduced MTHFR function associated with common genetic variants (e.g., the 677C>T polymorphism) can compromise this conversion efficiency and has been linked to elevated homocysteine and impaired folate status in susceptible individuals[27]. Collectively, these considerations suggest that probiotic-mediated delivery of 5-MeTHF may offer advantages over folic acid supplementation, providing a potentially more efficient and physiologically relevant strategy to improve folate status *in vivo*.

Indeed, animal experiment showed supplementation with the probiotic cocktail improved folate levels and reduced Hcy, but these effects were significant only in the high dose group. This dose dependence aligns with the qPCR findings showing that only high-dose probiotic administration resulted in an obvious increase in the relative abundance of supplemented strains in feces. Probiotics are defined as live microorganisms that, when administered in adequate amounts, confer a health benefit on the host[28]. Similarly, our data also suggest the existence of a threshold effect: a sufficient inoculum is required to achieve meaningful gut exposure (or transient persistence) and thereby elicit measurable improvements in folate availability and downstream one-carbon metabolism reflected by reduced Hcy. Therefore, future clinical studies are warranted to investigate the precise effective dose in human subjects.

Probiotic supplementation has emerged as a promising strategy to modulate the composition and functional profile of the gut microbiota[29]. Indeed, the probiotic cocktail increased microbial alpha diversity (Shannon index) and partially mitigated the compositional shifts induced by a folate-deficient diet. These community-level changes provide a plausible indirect mechanism whereby probiotics may enhance host folate status not only through direct folate vitamer production, but also by restoring or supporting a microbiota configuration enriched in folate-related functions. Genome-based predictions suggested that a substantial fraction of resident gut commensals may be capable of folate biosynthesis, particularly at steps related to pABA supply[25, 30]. Consequently, the improvement in folate status and Hcy metabolism observed here may result from (i) direct production of bioactive folate by the administered strains, (ii) microbiome restructuring that enhances endogenous folate-producing capacity, or (iii) a combination of both via cross-feeding and functional network effects. Dissecting these contributions will require additional studies, such as isotope-tracing to track microbially derived folate into host pools, gnotobiotic or defined-consortium models to test strain-strain interactions, and targeted depletion/reconstitution approaches to isolate direct versus community-mediated mechanisms.

## Conclusion

In summary, this work provides mechanistic and functional evidence that selected probiotic strains can produce bioactive folate 5-MeTHF, and that supplementation improves host folate status and lowers homocysteine *in vivo*. The data indicate that probiotic effects likely depend on both strain-specific metabolic capacity and precursor cross-feeding and gut community restoration under folate deficiency. These findings underscore the potential of high-folate-producing probiotics serve as a safer, more physiological intervention for folate deficiency and hyperhomocysteinemia, particularly in individuals carrying MTHFR polymorphisms.

## Supporting information

Graphic abstract

## Ethics approval and consent to participate

The study protocol, including the collection and analysis of human stool samples, was approved by the Ethics Committee of the Institute of Biomedical Sciences, Anhui Medical University (Approval No.: CH1095). All experimental methods were performed in accordance with the relevant guidelines and regulations. Informed consent was obtained from all participants prior to sample collection.

## Consent for publication

All authors consented to the publication of the manuscript.

## Authors’ contributions

Mingfang PAN: Methodology, Software, Formal Analysis, Investigation, Data Curation, Writing - Original Draft. Changming YE: Resources, Project Administration. Yun SON: Writing - Review & Editing. Minqing TIAN: Resources, Project Administration. Runming WANG: Conceptualization, Supervision, Writing - Review & Editing. Ping CHEN: Conceptualization, Supervision, Project Administration, Funding Acquisition, Writing - Review & Editing. All authors have read and approved the final manuscript.

## ACKNOWLEDGMENTS

This work was supported by the Science and Technology Major Project of the Shenzhen Municipal Science and Technology Innovation Committee (Grant No. KJZD20230923114408018): R& D of Gut Microbiota-Based Probiotic for Stroke Prevention and the funding from the Development and Reform Commission of Shenzhen Municipality(grant no.XMHT20240104002). The funding agency had no involvement in study design, data interpretation, or manuscript preparation.

