## Supplementary table and figure for "Isolation of folate-producing probiotics and its regulatory effects on homocysteine metabolism and gut microbiota composition"

**Supplementary Table 1. Quantitative PCR (qPCR) primers designed for detection of administered probiotic species**

| Targeted probiotic species | Primer name | Primer(5’→3’) | Reference |
|---|---|---|---|
| *Weizmannia coagulans* | B.coa-F | CGAACTCGAAGAATATGATGACA | [1] |
| | B.coa-R | TCATCTTTCGACATGATTTGG | |
| *Lactiplantibacillus plantarum* | LP-F | CAAGCCTACGCAACTATCAACC | [2] |
| | LP-R | ACCGTGGGCAAACATTTTATT | |
| *Bifidobacterium breve* | BiBRE-F | CCGGATGCTCCATCACAC | [3] |
| | BiBRE-R | ACAAAGTGCCTTGCTCCCT | |
| *Bifidobacterium longum* | BIL-F | GTTCCCGACGGTCGTAGAG | [4] |
| | BIL-R | GTGAGTTCCCGGCATAATCC | |
| *Bifidobacterium animalis* | B.ani-F | GAAAAGGTTGAGAAGGACTTCAACC | [4] |
| | B.ani-R | GAAGCCCTCACCCTCAG | |

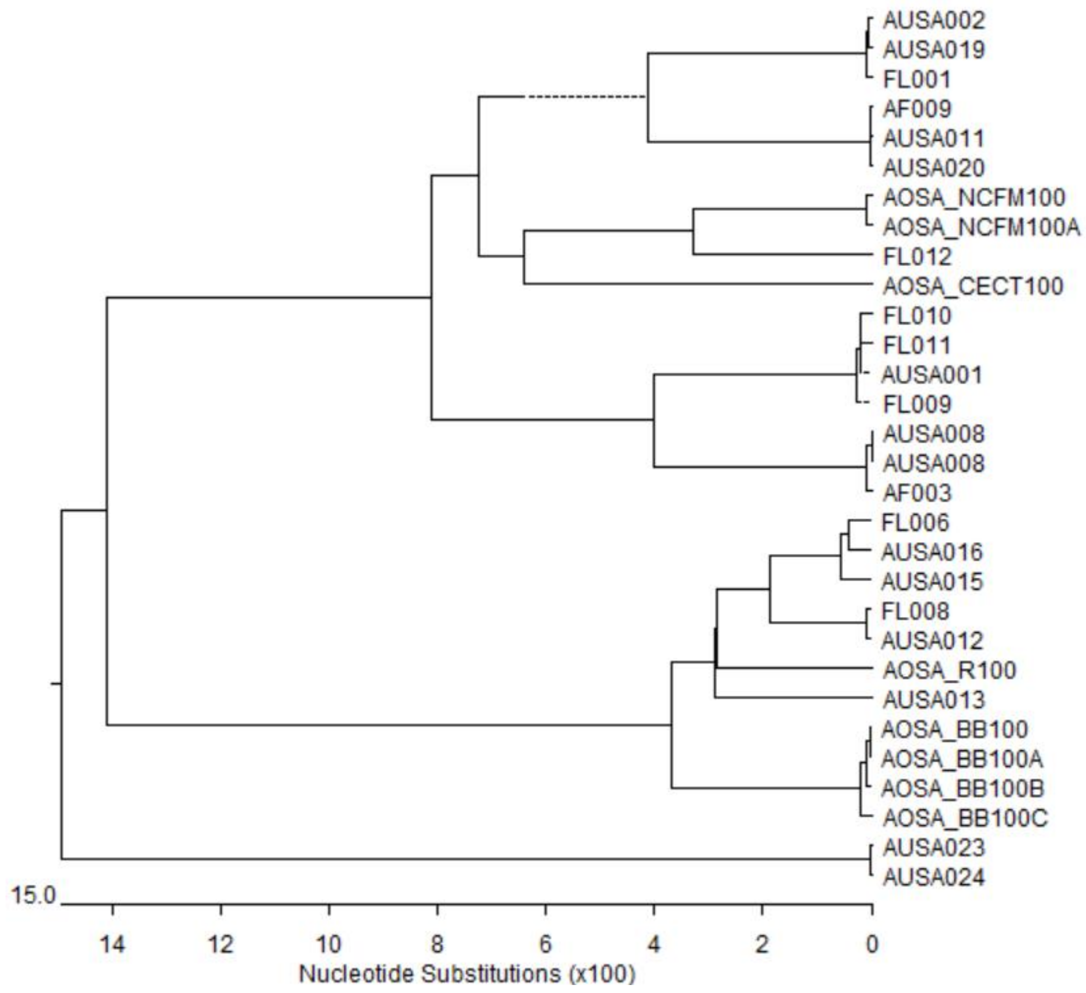


**Supplementary Figure 1.** Phylogenetic tree of the isolated probiotic strains based on 16s rRNA gene sequence.

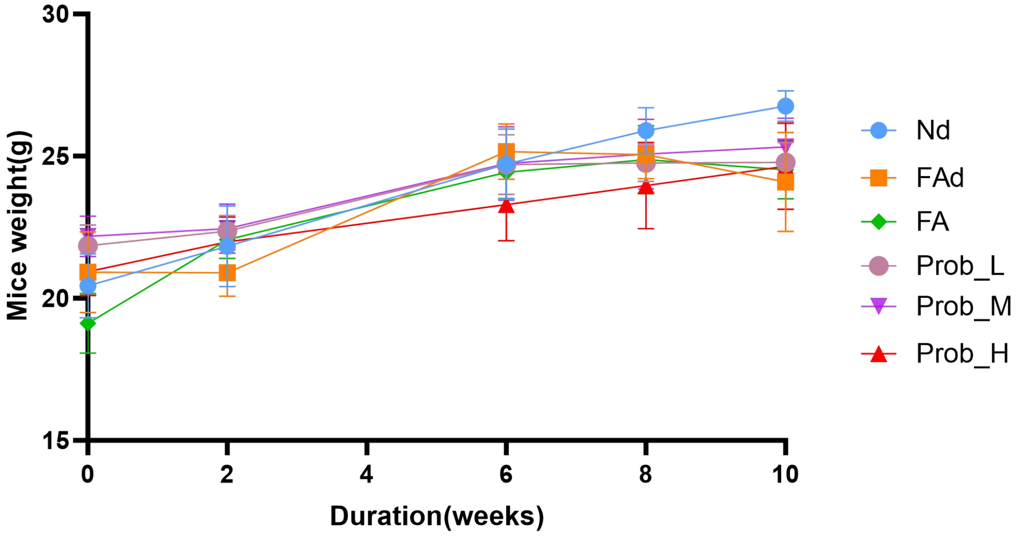


**Supplementary Figure 2.** The weight change of the mice during intervation. Nd: Normal diet/PBS; FAd: Folate-deficient diet/PBS; Prob_H: Probiotic (high dose, 1010CFU); Prob_M: Probiotic(medium dose, 109CFU); Prob_L: Probiotic (low dose, 108CFU); FA: Folic acid (73ug/kg) .
