## Supplementary figures and images for "Isolation of folate-producing probiotics and its regulatory effects on homocysteine metabolism and gut microbiota composition"

### Graphic abstract

Graphical Abstract:

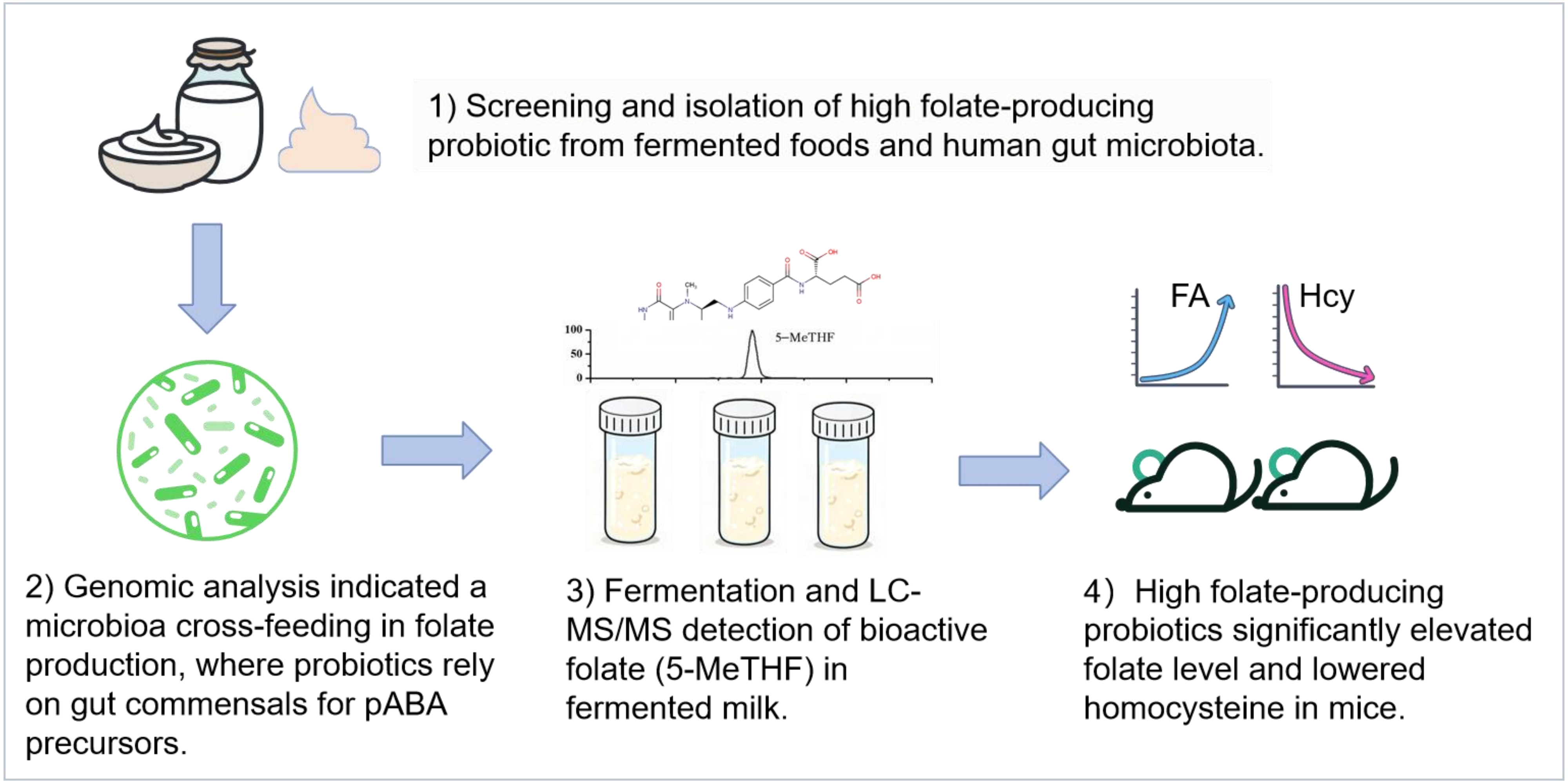
